# New mega dataset combined with deep neural network makes a progress in predicting impact of mutation on protein stability

**DOI:** 10.1101/2022.12.31.522396

**Authors:** Marina A Pak, Nikita V Dovidchenko, Satyarth Mishra Sharma, Dmitry N Ivankov

**Author notes:** Equal contribution (Marina A Pak, Nikita V Dovidchenko, Satyarth Mishra Sharma).

## Abstract

Prediction of proteins stability change (ΔΔG) due to single mutation is important for biotechnology, medicine, and our understanding of physics underlying protein folding. Despite the recent tremendous success in 3D protein structure prediction, the apparently simpler problem of predicting the effect of mutations on protein stability has been hampered by the low amount of experimental data. With the recent high-throughput measurements of mutational effects in ‘mega’ experiment for ~850,000 mutations [Tsuboyama et al., bioRxiv, 2022] it becomes possible to apply the state-of-the-art deep learning methods. Here we explore the ability of ESM2 deep neural network architecture with added Light Attention mechanism to predict the change of protein stability due to single mutations. The resulting method ABYSSAL predicts well the data from the ‘mega’ experiment (Pearson correlation 0.85) while the prediction of ΔΔG values from previous experiments is more modest (Pearson correlation 0.50). ABYSSAL also shows a perfect satisfaction of the antisymmetry property. The ABYSSAL training demonstrated that the dataset should contain around ~100,000 data points for taking advantage of the state-of-the-art deep learning methods. Overall, our study shows great perspectives for developing the deep learning ΔΔG predictors.

## Introduction

Prediction of protein stability change (ΔΔG) upon mutation is one of the most important unsolved problems of structural bioinformatics (Toplak et al., 2021; Kalman et al., 2020; Wang et al., 2020; Pancotti et al., 2022; Pak & Ivankov, 2022). The recent success of AlphaFold in predicting 3D protein structure at near-to-experimental accuracy showed the perspectives of deep learning techniques for solving biological problems (Jumper et al., 2021). The vast amount of known protein sequences (Uniprot Consortium, 2012) and known crystallographic structures (Berman et al., 2000) played a crucial role in AlphaFold’s success. Field of ΔΔG prediction always suffered from the lack of data: by the middle of the year 2022 only ~14k experimental records were collected (Xavier et al., 2021) which may be too low to learn the ΔΔG prediction by a deep neural network.

Recently, Tsuboyama et al. published the experimentally measured ΔΔG values for 851,552 mutations, with 376,918 of them being high-quality single mutations (Tsuboyama et al., 2022). The dataset is much larger than any dataset used before and has no bias towards ‘truncating’ mutations to smaller amino acids, especially to alanine. Thus, it provides a unique opportunity to develop an unbiased state-of-the-art ΔΔG predictor using one of the powerful deep learning models developed recently (Lin et al., 2022).

Here we present ABYSSAL (**Mega** dataset and **Deep** neural network with **a**ttention-**l**ike mechanism), the first predictor of protein stability change due to single mutation trained on such a big amount of data. ABYSSAL takes advantage of the state-of-the-art deep neural network model ESM2 (Lin et al., 2022). ABYSSAL predicts experimental ΔΔG values at the level of Pearson correlation coefficient (PCC) equal to 0.85, which amounts to near-to-experimental quality (Tsuboyama et al., 2022; Potapov et al., 2009; Xavier et al., 2021). We have shown that a training dataset should contain around ~100,000 data points is enough to take full advantage of the current state-of-the-art deep neural network models like ESM2 (Lin et al., 2022).

## Materials and Methods

### Dataset

We took the experimental data on protein stability changes (ΔΔG) upon mutations from (Tsuboyama et al., 2022) where ΔΔG values were estimated from cleavages by proteases. We downloaded the file “K50_dG_Dataset1_Dataset2.csv” from the Zenodo repository https://zenodo.org/record/7401275#.Y6st59JBx Dassociated with the paper (Tsuboyama et al., 2022). Out of 851,552 stability change data for 542 reference proteins, we removed records (i) having tag ‘unreliable’, (ii) when no mutation was introduced, (iii) associated with insertions and/or deletions, (iv) associated with multiple mutations. The filtered dataset contained data on 376,918 single mutations from 396 proteins. We reconstructed the sequences of the wild type proteins from the mutated sequence in the “aa_seq” column and the mutation in the column “mut_type” of the original dataset. The effect of mutation from the column “ddG_ML” was multiplied by −1 to convert the values into folding free energy changes (negative values denote stabilization). We called the resulting filtered and processed dataset Mega dataset.

### Mega dataset split into training, testing, and validation datasets

We used two approaches for splitting the Mega dataset into training, testing, and validation sets. The first approach is based on protein sequence identity cutoffs. Firstly, we performed all-against-all protein BLAST (Altschul et al., 1997) of protein sequences of the Mega dataset with the E-value of 10^-5^. The smallest sequence identity cutoff we could use to have sufficient data in the test and validation sets was 35%. We split the data in such a way that proteins from the testing and validation sets were similar to the proteins of the training set at maximum by 35% of sequence identity. The remaining 367,858 mutations comprised the training dataset. From the remaining 9,060 mutations aimed to test and validate the results we assigned randomly 6,043 (two thirds) to the test set while the other 3,017 mutations (one third) comprised the validation test. The resulting sets are denoted as MegaTrain, MegaTest and MegaValidation, respectively. The second approach is a straightforward naive approach – a random split into train and test by 0.8/0.2 ratio; it was used to explore sequence-identity-unaware DDG prediction. In addition we have trained our model on a popular training set S2648 (Dehouck et al., 2009).

The sizes and the number of proteins in the training, testing and validation sets are represented in Table S1. All data sets were symmetrized by adding the reverse mutations which doubled the sizes of every dataset.

### ΔΔG validation datasets

In addition to the MegaValidation dataset we used popular datasets of experimental ΔΔG data: p53 (Pires et al., 2014) (42 mutations), Myoglobin (Kepp, 2015) (134 mutations), and Ssym (Pucci et al., 2018) (342 mutations). We have also used a recently developed dataset S669 (Pancotti et al., 2022) (669 mutations). It includes curated data dissimilar at 25% of sequence identity to widely used training sets S2648 (Dehouck et al., 2009) and VariBench (Nair, Vihinen, 2012). From all validation datasets we excluded mutations in proteins that are similar to the proteins in Mega dataset at sequence identity cutoff of 25%. This resulted in 420 mutations comprising the filtered version of the S669 dataset. The description of validation datasets is presented in Table S2.

### Metrics

We used six metrics for assessment of ΔΔG prediction: the Pearson and Spearman correlation coefficients between true ΔΔG values and predicted values (PCC, SCC), mean standard error (MSE), the accuracy of predicting a class of mutation (ΔΔG < 0 – stabilizing, ΔΔG ≥ 0 – destabilizing), and antisymmetry metrics PCC(f-w) and the bias <δ>. PCC(f-w) is the Pearson correlation coefficient between the forward and the reverse mutations (Usmanova et al., 2018; Pancotti et al., 2022):

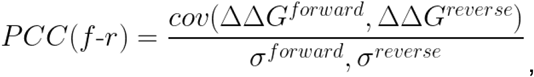

where cov is the covariance and σ is the standard deviation.

The bias <δ> is the average bias toward either forward or reverse mutations (Usmanova et al., 2018; Pancotti et al., 2022):

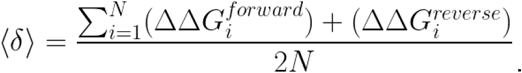

### Training procedure

Training on MegaDataset with random split was performed in two steps. At first step the model learned for 50 epochs with learning rate 10^-4^; after that the resultant model was fine-tuned for another 50 epochs with learning rate 10^-5^. For the split based on sequence identity cutoffs, training took place for 72 epochs with an initial learning rate of 10^-4^ decaying by a factor of 0.97 every epoch.

### Neural network architecture

#### Embeddings

To provide sequence representations we have used an ESM2 (esm2_t33_650M_UR50D) model (Lin et al., 2022) which was shown to be a state-of-the-art solution in a number of sequence analysis-related tasks (Lin et al., 2022). In current analysis, we have used two embeddings per sequence: one for the original sequence and one for the mutated one. Moreover, to underline the effect of a given position and given amino acid mutation, instead of whole sequence representation, embeddings for the specific token (amino acid) both for the original amino acid and mutated one from the last ESM2 layer were extracted.

#### ABYSSAL general description

The proposed model (Fig 1c) represents a variation of the Siamese network types of networks (Bromley et al., 1993). It has two inputs which are preprocessed inside the model (see Light Attention trick section and Fig 1a) and then both outputs are concatenated. We believe that the concatenation of both outputs in combination with providing both directions of mutations within the dataset (original amino acid → mutated amino acid and mutated amino acid → original amino acid within the train dataset) essentially helps the model to learn the physical property of antisymmetry of ΔΔG. Obtained concatenated vector then is fed into several fully connected layers (Fig 1b) and output of the last layer is the predicted ΔΔG (Fig 1c).

**Figure 1.**
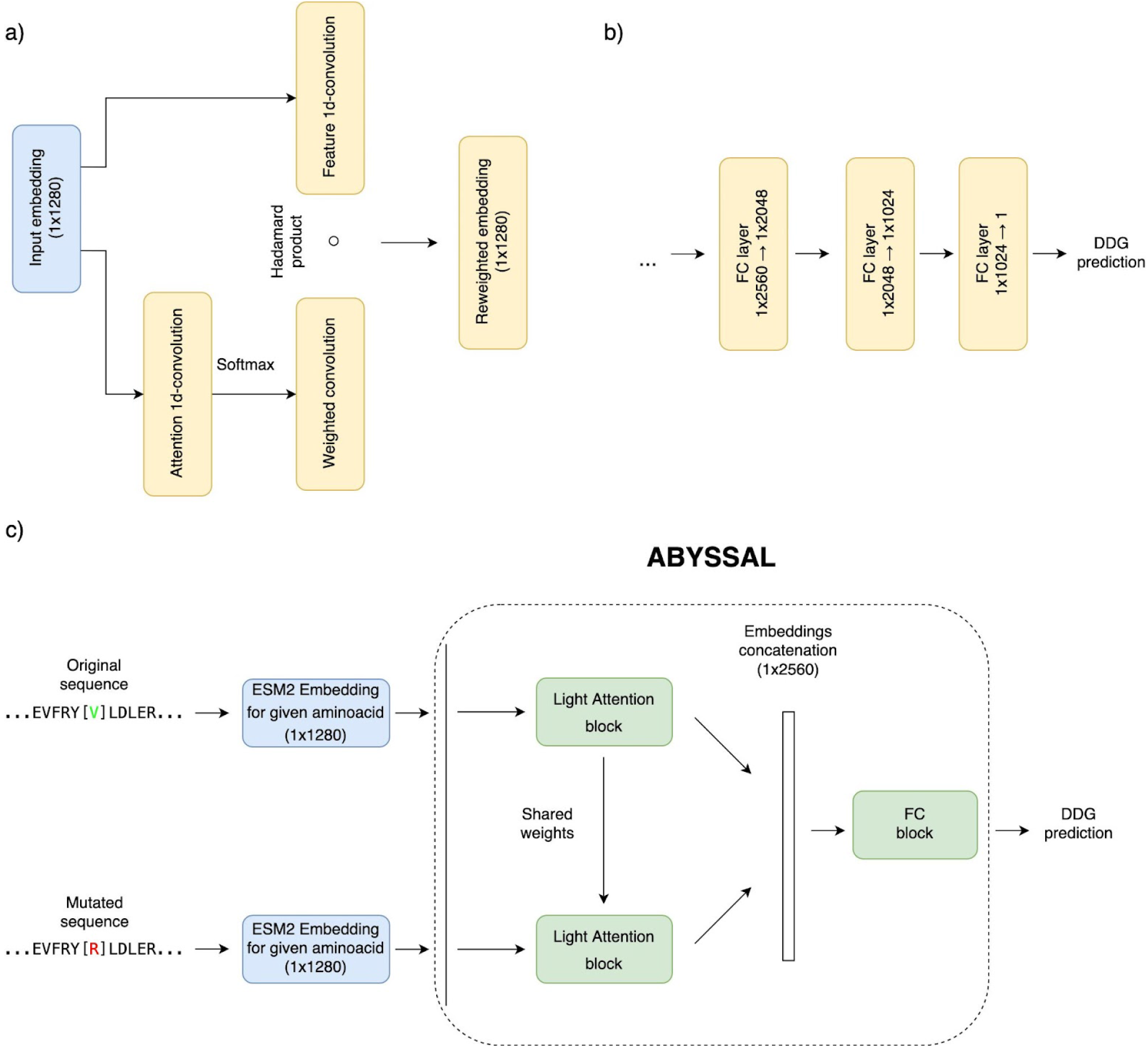
General scheme of the ABYSSAL: a) scheme of Light Attention (Stärk et al., 2021) block; b) scheme of fully connected (FC) block which consists of several fully connected layers; c) general scheme of ABYSSAL and flow of data: at first step we obtain embeddings for both mutated and original amino acid by using ESM2 [Language models of protein sequences at the scale of evolution enable accurate structure prediction] general language model, than both embeddings do serve as an input into ABYSSAL. Both embeddings are preprocessed through Light Attention block (subfigure “a”) in a siamese network fashion. Preprocessed by the Light Attention block, both embeddings are concatenated and fed in a FC block (subfigure “b”) which represents a set of several fully connected layers which outputs a predicted ddg.

#### Light Attention trick

It is usually a good decision to allow a network to reweight incoming embedding vectors so the relevant data could gain more attention and thus the network would converge better. Light attention (Stärk et al., 2021) is one of the proposed tricks which was shown to be helpful in such tasks; therefore, we incorporated it into the ABYSSAL. In the Light Attention trick (fig 1a) two vectors of the same shape are created by two 1d convolutions (kernel size was set to 9, stride to 1 and padding was set to 4). One of the obtained vectors serves as an inner model embedding representation and the other (by applying softmax) helps to reweight obtained representation in a attention-like manner. Both vectors are then multiplied in an element-wise manner (Hadamard product).

## Results and Discussion

### ABYSSAL learning design

The aim of our study is to explore perspectives of deep learning to the problem of predicting change of protein stability (ΔΔG) upon single mutation in globular proteins, the conceptually simplest task of protein design. For that, we developed ABYSSAL, a method for ΔΔG prediction trained on the data from the recently published dataset (Tsuboyama et al., 2022). After filtering, the dataset contained 376,918 high-quality single mutations in 396 small single-domain globular proteins, see Methods. The big size of the dataset allows using the most advanced deep learning techniques, so we chose ESM2, a language model, which showed previously its usability in the protein structure prediction (Lin et al., 2021). Additionally, we used Light Attention mechanism which proved to be useful in previous biological research (Stärk et al., 2021).

To follow the best practices in machine learning, we divided the dataset into training set (MegaTrain), test set (MegaTest), and validation set (MegaValidation) as described in Methods. This is useful when the best model is selected out of many developed models based on the performance on the test set while the actual performance is reported for the validation set. Although we developed only one model, we split the Mega dataset into training, test, and validation sets for future investigations.

Another thing to address is the similarity between test/validation and training sets. Indeed, too high similarity of the test/validation sets with the training set may lead to overestimating the performance of the method and compromising the overall results. Obviously, one must avoid same mutations being present in the test/validation and the training sets. However, little is known about the similarity parameters between mutations to avoid overlearning of statistical and machine learning methods. To be on the safe side, we used the rules established for protein sequence-to-structure relationship where sequence identity higher than the twilight zone ~20-40% usually results in highly similar protein 3D structures. Specifically, we constructed the MegaTest and MegaValidation sets so that they contain proteins that are dissimilar to those from the MegaTrain set. At the same time, we have checked that the number of mutations in the MegaTest and MegaValidation sets was large enough to estimate reliably of the ABYSSAL performance. This resulted in the sequence identity threshold of 35%, i.e., any protein from the MegaTest and MegaValidation sets has the sequence identity less than 35% to any protein in the MegaTrain set (see Methods). After the split, MegaTrain set was represented by 367,858 mutations from 388 proteins; from the remaining 9060 mutations in 8 proteins 3017 were randomly chosen mutations and assigned to MegaValidation set, see Table S1. To understand the performance of the model in sequence-identity-unaware regime, we trained the model using naive straightforward approach of split in 0.8/0.2 proportion. This resulted in a significantly higher performance without overfitting. However, since it is unknown about similarities rules for mutation, we do not show these preliminary data.

One of the main pitfalls of constructing most of the ΔΔG prediction methods in the past was their violation of the antisymmetry property: the effects of the forward and reverse mutations must be exactly opposite to each other, i.e., their average sum <δ> must sum up to zero while the correlation between the forward and reverse subsets must be −1 (Fariselli et al., 2015; Usmanova et al., 2018; Pucci et al., 2018). To avoid antisymmetry property violation, we followed the best machine learning practice and symmetrized the MegaTrain, MegaTest, and MegaValidation set, i.e., for every mutation in a dataset we added the corresponding reverse mutation, which resulted in doubling the sizes of all sets (we name further these combined sets as ‘symmetric’). Additionally, we used special neural network architecture to complement model (introduced by dataset modifications) to antisymmetry property (see Methods) in a manner similar to (Benevenuta et al., 2021).

### Performance of ABYSSAL

We trained ESM2 model (Lin et al., 2021) with Light Attention trick (Stärk et al., 2021) on the MegaTrain dataset as described in Methods. The performance of the model on MegaTest set is shown in Table 1. The PCC was 0.85 which approaches the agreement to PCC = 0.91 between trypsin- and chymotrypsin-based data from the Mega dataset (Tsuboyama et al., 2022). For the MegaValidation dataset ABYSSAL showed the same PCC = 0.85, see upper panel of Fig. 2. The antisymmetry property was satisfied perfectly, with PCC between forward and reverse mutations being −0.98 and <δ> of just 0.01 kcal/mol.

**Table 1.** Performance metrics of ABYSSAL on MegaTest.

| Metrics | Symmetric | Forward | Reverse | Antisymmetry |  |
| --- | --- | --- | --- | --- | --- |
| | | | | PCC(f-w) | $\langle\delta\rangle$ |
| PCC | 0.85±0.00 | 0.77±0.01 | 0.77±0.01 | -0.98 | -0.01 |
| SCC | 0.81±0.01 | 0.73±0.01 | 0.73±0.01 |  |  |
| MSE, kcal/mol | 0.89 | 0.89 | 0.88 |  |  |
| Accuracy | 0.79 | 0.79 | 0.79 |  |  |

**Figure 2.**
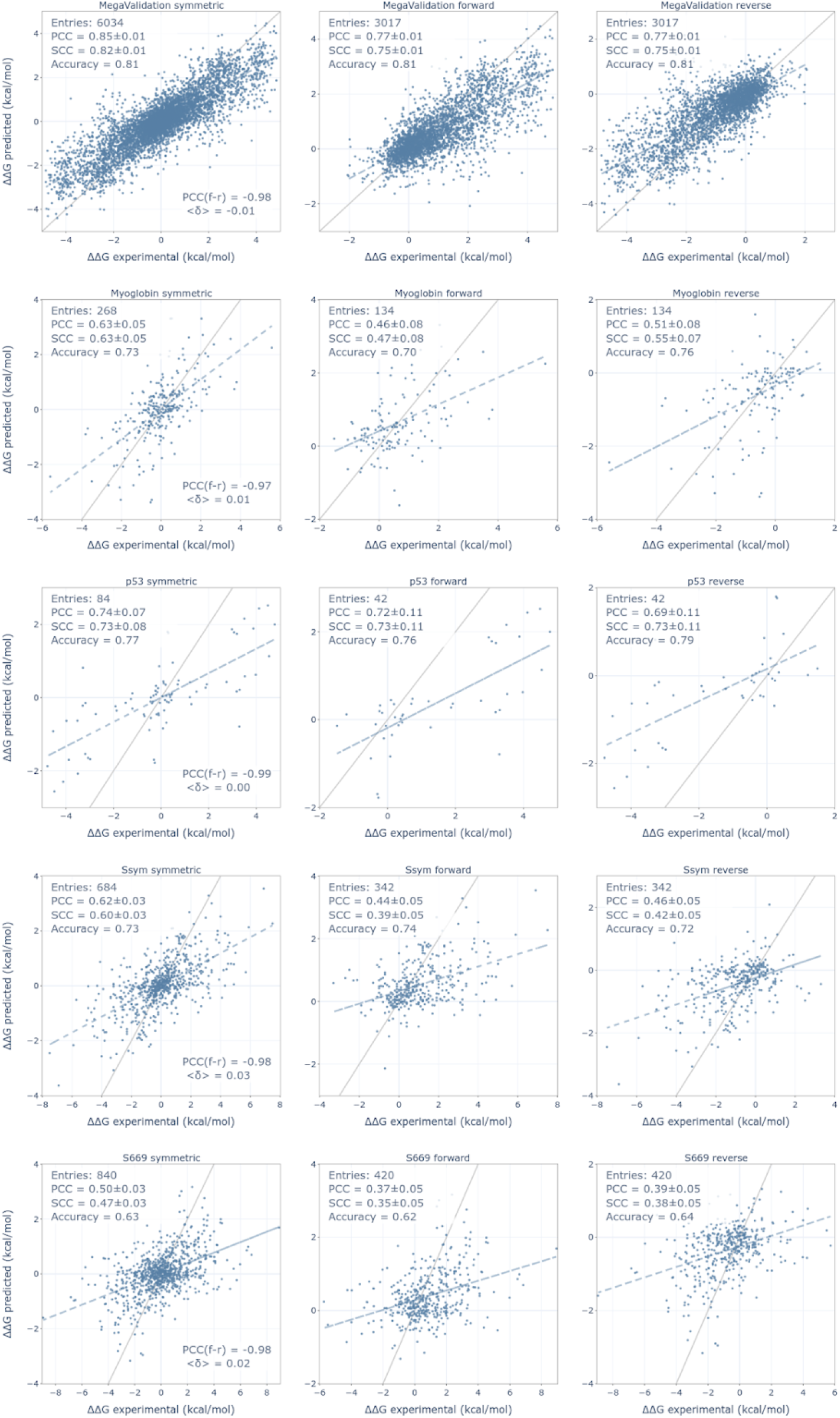
Performance on validation sets (from top to bottom): for MegaValidation, Myoglobin, p53, Ssym, and S669 datasets. In every row the results are given for full dataset containing both forward and reverse mutations (left), for forward mutations only (middle), for reverse mutations only (right). The following metrics are given in the plots: number of points (‘entries’), PCC, SCC, Accuracy and asymmetry metrics (PCC between forward and reverse mutations and average bias <δ>) for full dataset. Line y = x is drawn in gray. The best fit is drawn as dashed line. Note that S669 dataset was filtered for mutations in proteins homologous to that in proteins from the training dataset of ABYSSAL so it contains 420 mutations. See text for details.

It should be noted that in the Mega paper ΔΔG values were estimated from proteins’ propensities to be cleaved by trypsin/chemotrypsin proteases. The agreement between the ΔΔG estimations and previous ΔΔG values from literature was in the range of PCC between 0.72 and 0.96 (Tsuboyama et al., 2022), which is very high but not ideally perfect. This means that ΔΔG estimations from the Mega paper appear as close proxies of the actual ΔΔG values. Since the declared problem of structural bioinformatics is the prediction of ΔΔG values, we are interested in how our ABYSSAL method performs on the previously measured ΔΔG values. We therefore assessed the ABYSSAL’s performance on the ΔΔG datasets ‘p53’ (Pires et al., 2014), ‘myoglobin’ (Kepp, 2015), ‘ssym’ (Pucci et al., 2018), and ‘s669’ (filtered versions, see below) (Pancotti et al., 2022) previously widely used in literature. Since the ΔΔG data from previous publications were not used for training ABYSSAL, the datasets act as a completely independent check of the ABYSSAL’s ability to predict the ΔΔG values. The performance of ABYSSAL is given in Fig.2, lower panels.

To compare the ABYSSAL performance with that of previously developed ΔΔG predictors we chose the biggest dataset available for such comparison, the recently combined dataset ‘s669’ (Pancotti et al, 2022). To avoid testing of ΔΔG predictors on the mutations in homologous proteins presented in the training sets of ΔΔG predictors we removed from ‘s669’ dataset mutations that were part of any ΔΔG predictor’s train set. It turned out that the arbitrary chosen sequence identity cut-off of 25% retains 420 out of 669 mutations (comprising 63% of the original ‘s669’ dataset). The results of the comparison are given in Table 2. ABYSSAL and INPS-Seq demonstrated the best performance of PCC = 0.50; ABYSSAL, INPS-Seq, and ACDC-NN showed best MSE of 1.74 kcal/mol. As for other metrics, SCC, accuracy, and antisymmetry, ABYSSAL was consistently among the best 4-5 predictors. For smaller datasets (Myoglobin, p53, and ssym) ABYSSAL performed sometimes better, sometimes worse than the other methods (data not shown).

**Table 2.**
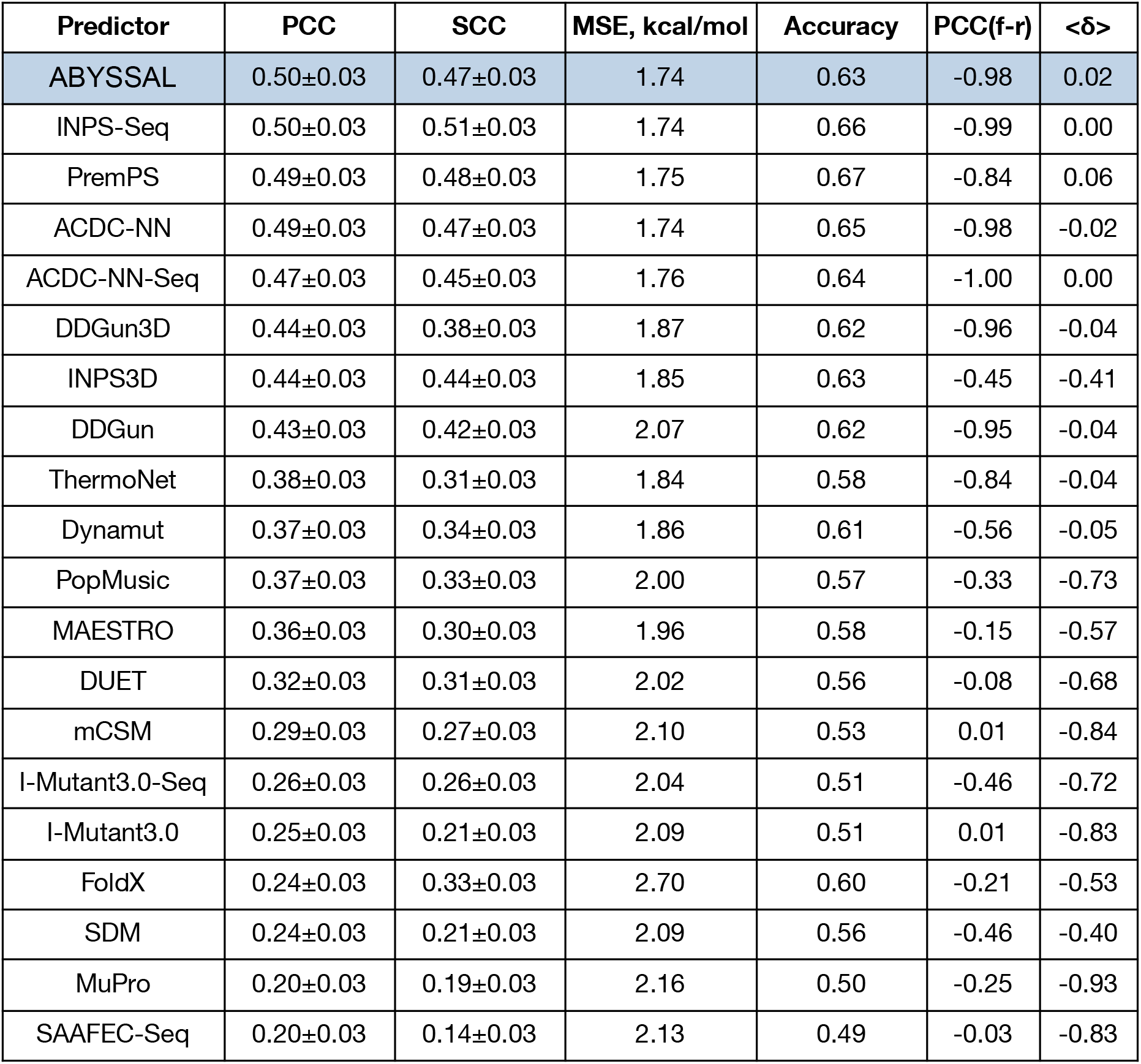
Performance of ΔΔG predictors on S669 dataset.

We were surprised that ABYSSAL did not show significant improvement on the S669 dataset. We did not perform a thorough study of that result yet; however, we can speculate and the reasons might be following. ABYSSAL was trained to predict ΔΔG estimates measured from cleavage by proteases (‘proxies’ to actual ΔΔG values), see PCC = 0.85 above. On the contrary, the other methods do predict the actual ΔΔG values; the S669 dataset contains the actual ΔΔG values, too.

The values in Table 1 were calculated from data presented in (Pancotti et al., 2022). The data are symmetrical and sorted by PCC in descending order.

### Influence of training set size on the ABYSSAL performance

One of the known limitations of neural networks is that they require a lot of data for their training. Thank to the Mega dataset containing 376,918 mutations we could train the ABYSSAL on 367,858 single mutations to near-to-experimental accuracy of the ΔΔG-estimates-producing method (Tsuboyama et al., 2022). However, we wonder about the minimal dataset size for training the deep learning model we chose here. To investigate the effect of the training dataset size, we repeated the learning of our model using smaller subsets of the original training dataset covering the wide range of data points from 2,441 to 367,858. The number 2,441 was chosen because it is close to the number of mutations in the dataset S2648 containing 2648 points, which was very widely used for developing DDG predictors.

The resulting dependence of model’s performance on the MegaValidation dataset and ΔΔG validation datasets are given in Fig.3. We see that the performance on the MegaValidation dataset clearly improves with the size reaching plateau value of 0.85 after 87,773 dataset size. This means that at 35% sequence identity cut-off our model successfully extracted all possible information already from ‘only’ 87,773 (24% of the MegaTrain dataset) mutations. Importantly, the model trained on the dataset size of 2,441 shows good (PCC = 0.80) but statistically significantly lower performance. This should be kept in mind when trying to learn state-of-the-art deep learning models on datasets containing low number of mutations.

**Figure 3.**
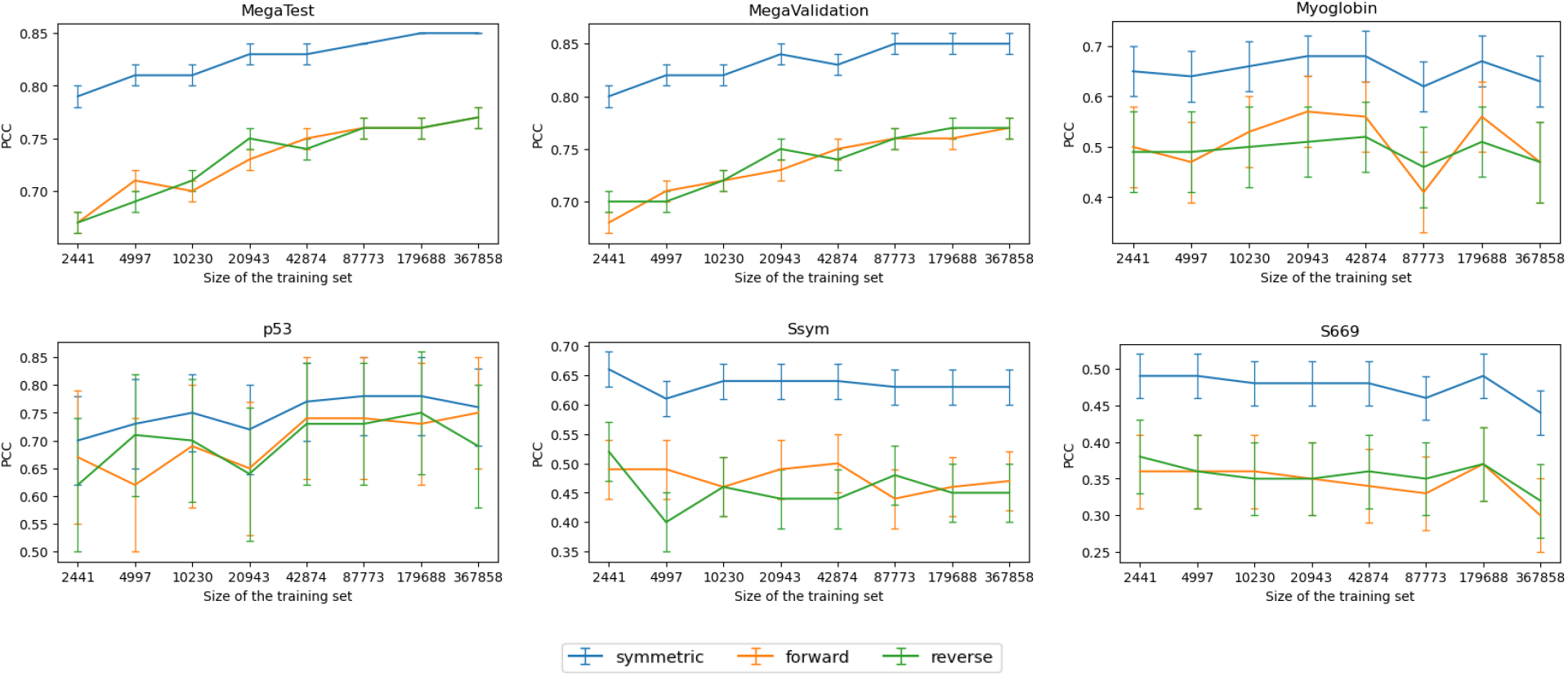
Dependence of the model performance as a function of the training dataset size.

Interestingly, the performance of ΔΔG predictions in other datasets showed completely random behavior. This could be an extra sign that actual ΔΔG values are of different nature from the ΔΔG estimates obtained in the Mega experiment (Tsuboyama et al., 2022). However, the small sizes of the independent ΔΔG datasets do not allow to make any statistically justified conclusions (Fig.3).

### S2648

To explore the perspectives of training our model on small datasets for the problem of ΔΔG prediction we trained our model directly on the S2648 dataset. We then made the cross-validation of the model’s performance on the MegaValidation dataset, and validations for the non-homologous part of the S669 dataset containing 411 data points, see Fig.4. We see that the trained model, indeed, is performing worse on the MegaValidation data (PCC = 0.71 compared to PCC = 0.80, see above), which confirms the slight intrinsic differences between ΔΔG estimates measured in the Mega dataset and actual ΔΔG values measured directly by biophysical methods. We also see that the trained model shows on S669 dataset lower but statistically indistinguishable correlation from ABYSSAL (0.48±0.03 vs. 0.50±0.03, see Table 2, p-value of Fisher r-to-z test 0.31). This may mean that if the amount of actual ΔΔG values would be higher, our model might achieve much better performance (compare with the training dataset size dependence in Fig.4).

**Figure 4.**
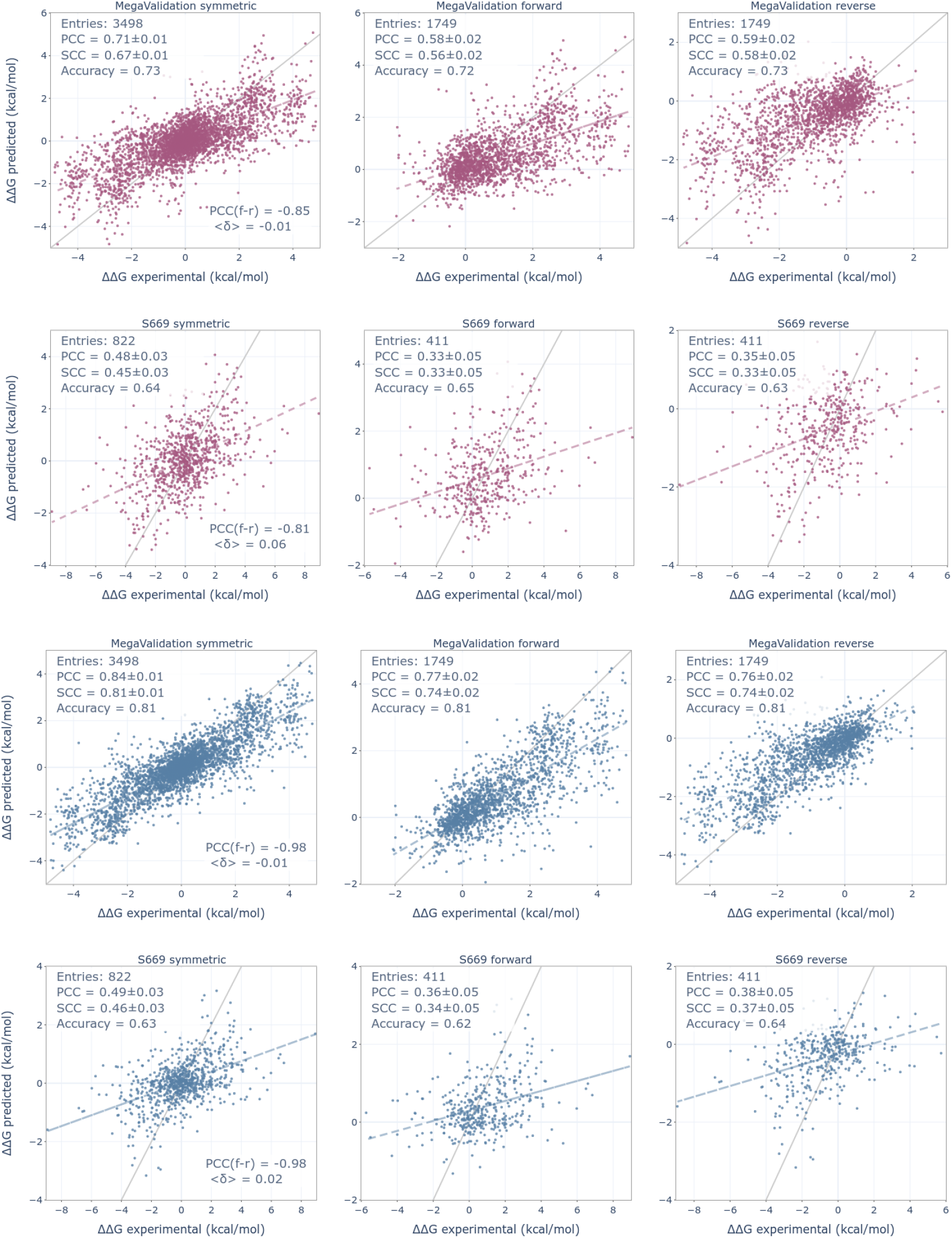
Performance of the model trained on S2648 (pink) dataset and performance of the model trained on MegaTrain (blue) dataset. Same subsets of validation datasets.

In the present study we explored perspectives of the training state-of-the-art deep learning methods We developed the method ABYSSAL which shows the near-to-experimental performance for ΔΔG prediction (PCC = 0.85). On the previous data ABYSSAL shows lower correlation (PCC = 0.50). We showed that around ~100,000 data points for taking advantage of the state-of-the-art deep learning methods. Overall, our study shows great perspectives for developing the deep learning ΔΔG predictors.

## Acknowledgements

The authors thank Alexander Medvedev for fruitful discussions. The authors acknowledge the usage of the Skoltech HPC cluster Zhores (Zakharov et al., 2019) for obtaining the results presented in this paper.

## Supplementary Tables

**Table S1.** Training and testing datasets used in the study. The values stated for the dataset before symmetrization.

| Dataset name | Size | Number of proteins | Portion of the Mega dataset |
| --- | --- | --- | --- |
| MegaTrain | 367858 | 388 | 0.980 |
| MegaTest | 6043 | 8 | 0.016 |

**Table S2.** Validation datasets used in the study. The values stated for the dataset before symmetrization.

| Name | Original |  | 25% cutoff with Mega dataset |  | 25% cutoff with Mega dataset and S2648 |  |
| --- | --- | --- | --- | --- | --- | --- |
|  | Size | Number of proteins | Size | Number of proteins | Size | Number of proteins |
| MegaValidation | 3017 | 8 | 3017 | 8 | 1749 | 5 |
| Myoglobin | 134 | 1 | 134 | 1 | 0 | 0 |
| p53 | 42 | 1 | 42 | 1 | 0 | 0 |
| Ssym | 342 | 15 | 342 | 15 | 0 | 0 |
| S669 | 669 | 94 | 420 | 86 | 411 | 84 |

